# Native plasmid-encoded mercury resistance genes are functional and demonstrate natural transformation in environmental bacterial isolates

**DOI:** 10.1101/764415

**Authors:** Ankita Kothari, Drishti Soneja, Albert Tang, Hans Carlson, Adam M. Deutschbauer, Aindrila Mukhopadhyay

**Author notes:** Address correspondence to Aindrila Mukhopadhyay,.

## Abstract

Plasmid-mediated horizontal gene transfer (HGT) is a major driver of genetic diversity in bacteria. We experimentally validated the function of a putative mercury resistance operon present on an abundant 8 kilobase pair native plasmid found in groundwater samples without detectable levels of mercury. Phylogenetic analyses of the plasmid-encoded mercury reductases from the studied groundwater site show them to be distinct from those reported in proximal metal-contaminated sites. We synthesized the entire native plasmid and demonstrated that the plasmid was sufficient to confer functional mercury resistance in *Escherichia coli*. Given the possibility that natural transformation is a prevalent HGT mechanism in the low cell density environments of groundwaters, we also assayed bacterial strains from this environment for competence. We used the native plasmid-encoded metal resistance to design a screen and identified 17 strains positive for natural transformation. We selected 2 of the positive strains along with a model bacterium to fully confirm HGT via natural transformation. From an ecological perspective, the role of the native plasmid population in providing advantageous traits combined with the microbiome’s capacity to take up environmental DNA enables rapid adaptation to environmental stresses.

**Importance:** Horizontal transfer of mobile genetic elements via natural transformation has been poorly understood in environmental microbes. Here, we confirm the functionality of a native plasmid-encoded mercury resistance operon in a model microbe and then query for the dissemination of this resistance trait via natural transformation into environmental bacterial isolates. We identify seventeen strains including Gram-positive and Gram-negative bacteria to be naturally competent. These strains were able to successfully take up the plasmid DNA and obtain a clear growth advantage in the presence of mercury. Our study provides important insights into gene dissemination via natural transformation enabling rapid adaptation to dynamic stresses in groundwater environments.

## Introduction

Horizontal gene transfer (HGT) is the lateral movement of genetic material between cells (1) and has received increased attention due to the rapid emergence of multi-drug resistant bacteria (2, 3). HGT enables the incorporation of exogenous DNA to obtain new virulence and resistance traits. Plasmids are extrachromosomal entities that often confer novel and advantageous traits to the host. The plasmidome refers to the entire plasmid content of a microbial population, representing mobile genetic material that can be subjected to HGT. Plasmid-mediated HGT in bacteria can occur via conjugation (transfer of genetic material between bacterial cells by direct cell-to-cell contact), transduction (transfer of genetic material to a bacterium via a virus) and natural transformation (uptake of exogenously available genetic material by a bacterium) (4). Of these mechanisms, natural transformation is ostensibly the simplest, needing merely the presence of a competent bacterial cell to take up exogenously available DNA fragment(s). Natural transformation occurs when a cell is competent, a highly-regulated physiological state that reflects a window in which the proteins required for DNA binding, processing, and internalization are produced. The DNA taken up can be used as a nutrient source or a source of nucleotides for DNA synthesis, for genome repair, or to acquire variant alleles (5-8). Natural transformation has been detected in bacteria from all trophic and taxonomic groups including Archaebacteria suggesting that transformability evolved early in phylogeny (9). Thus, it has a significant impact on bacterial population dynamics as well as on bacterial evolution and speciation (9).

The Oak Ridge Field Research Center (ORFRC) (10-13) is a well-studied United States Department of Energy site that includes both areas with and without metal contamination, referred to as the contaminated and background sites, respectively. In a recent report (14) the plasmidome of the background site revealed the presence of a highly-abundant native plasmid containing putative mercury resistance genes (*mer*), despite a lack of detectable mercury contamination in the groundwater. It is not clear if the *mer* genes found at the background site are functional or how they came to be at that location. To get a deeper understanding, we examined the putative *mer*-encoding plasmid, p5343. We synthesized this plasmid and confirm that it provides functional mercury resistance in the model microbe *Escherichia coli*. We examined the phylogeny of the mercury reductases at ORFRC to obtain an understanding of its prevalence and distribution along with discussing the implications of our findings.

We use the synthesized the native plasmid to better understand its potential for HGT in a number of bacterial isolates from this environment. Since groundwater is a low cell density environment with fluctuating populations, we use natural transformation to examine the dissemination of the native plasmid-encoded metal resistance trait to bacterial isolates from the ORFRC site. Our results from these HGT assays are presented and reveal that natural transformation may play a critical role in the spread of resistance genes to divergent strains.

## Results

The lack of detectable mercury in the source groundwater for p5343 (14) raised the possibility that the *mer* genes on this plasmid are not functional. Vestigial *mer* operons (15, 16) have been reported and the closest MerA that has been experimentally confirmed to confer mercury resistance has only 47 % amino acid identity (17) to p5343 MerA (closest functional MerA identified based on PaperBLAST (18)). Therefore, to functionally examine the putative *mer* genes of p5343, *E. coli* was transformed with the *mer*-containing plasmid p5343_UC57 (Fig 1) or with a control plasmid pUC57 that lacks the *mer* genes. The transformed strains were assayed for mercury resistance at mercury chloride concentrations known to be inhibitory to *E. coli*. Consistent with a functional *mer* system, we observed higher mercury chloride half-maximal inhibitory concentration (IC_50_) values in *E. coli* carrying p5343_UC57 relative to the same strain with pUC57 (Fig 2). Specifically, at 14.3 μM mercury chloride, the strain with p5343_UC57 had a clear growth advantage. This is similar to the 10 μM mercury chloride concentration at which a growth advantage was reported earlier in *E. coli* transformed with mercury resistance genes (19). The proposed mercury reduction schematic based on previous illustrations (15, 20, 21), and SPOCTOPUS (22) predicted topology of *mer* operon in p5343, is depicted in Fig 3a. The p5343 only encodes genes providing narrow-spectrum mercury resistance (to inorganic mercury compounds like HgCl_2_ and Hg(NO_3_)_2_) while lacking genes encoding broad-spectrum mercury resistance (like methylmercury and phenyl mercuric acetate). When p5343_UC57 is cloned into *E. coli*, the genes *merP, merT* and *merF* likely aid in the transport of mercury while *merA* reduces Hg^+2^ to elemental mercury which volatilizes, leaving the bacterial environment mercury-free, resulting in improved mercury resistance. Thus, the native plasmid p5343 encodes a functional *mer* system that provides resistance to mercury.

**Fig 1:**
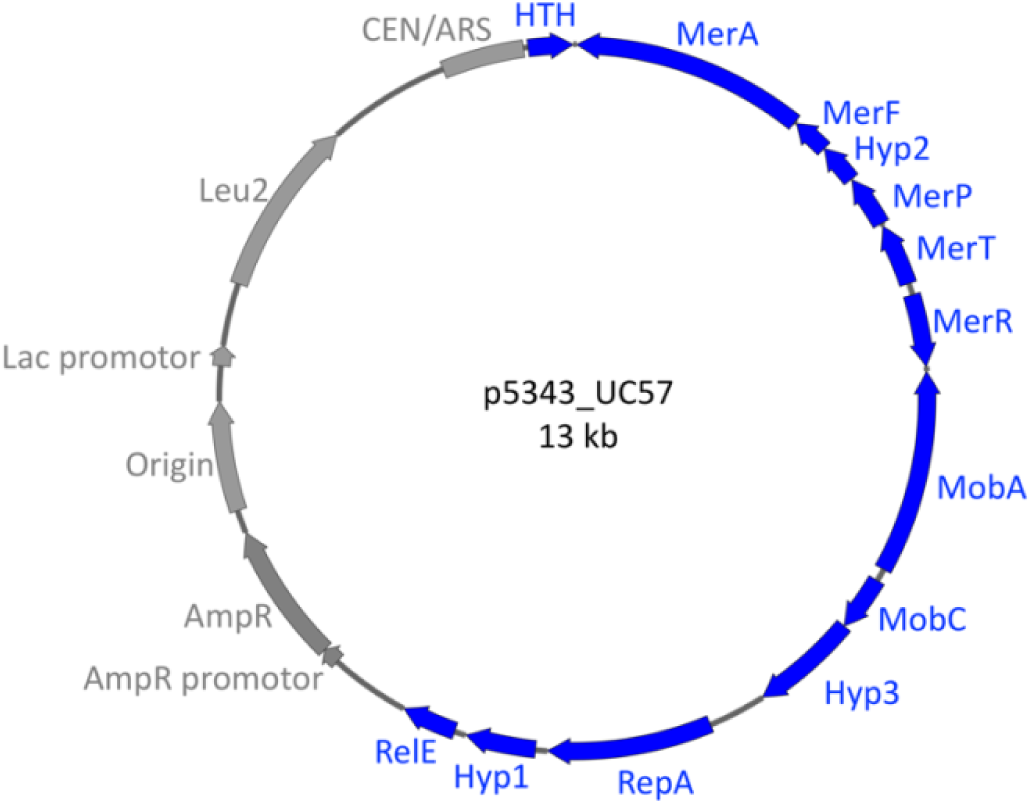
Plasmid map of p5343_UC57. The p5343 part (depicted in blue) encodes Mercuric ion reductase (MerA), Mercuric ion uptake protein (MerF), Hyp: Hypothetical protein (Hyp2), Mercuric transport protein (MerP), Mercuric transport protein (MerT), Regulator of Mercury resistance genes (MerR), Mobilization protein A (MobA), Mobilization protein C (MobC), Plasmid replication protein (RepA), Helix-turn-helix domain protein (HTH), RelE toxin (RelE), while the pUC57 part (depicted in grey) codes for promotor sequence for Ampicillin resistance gene (AmpR Promotor), Ampicillin resistance marker (AmpR), origin of replication, Lac promotor, LEU2 selection marker (Leu2) and elements ensuring plasmid maintenance (CEN/ARS).

**Fig 2:**
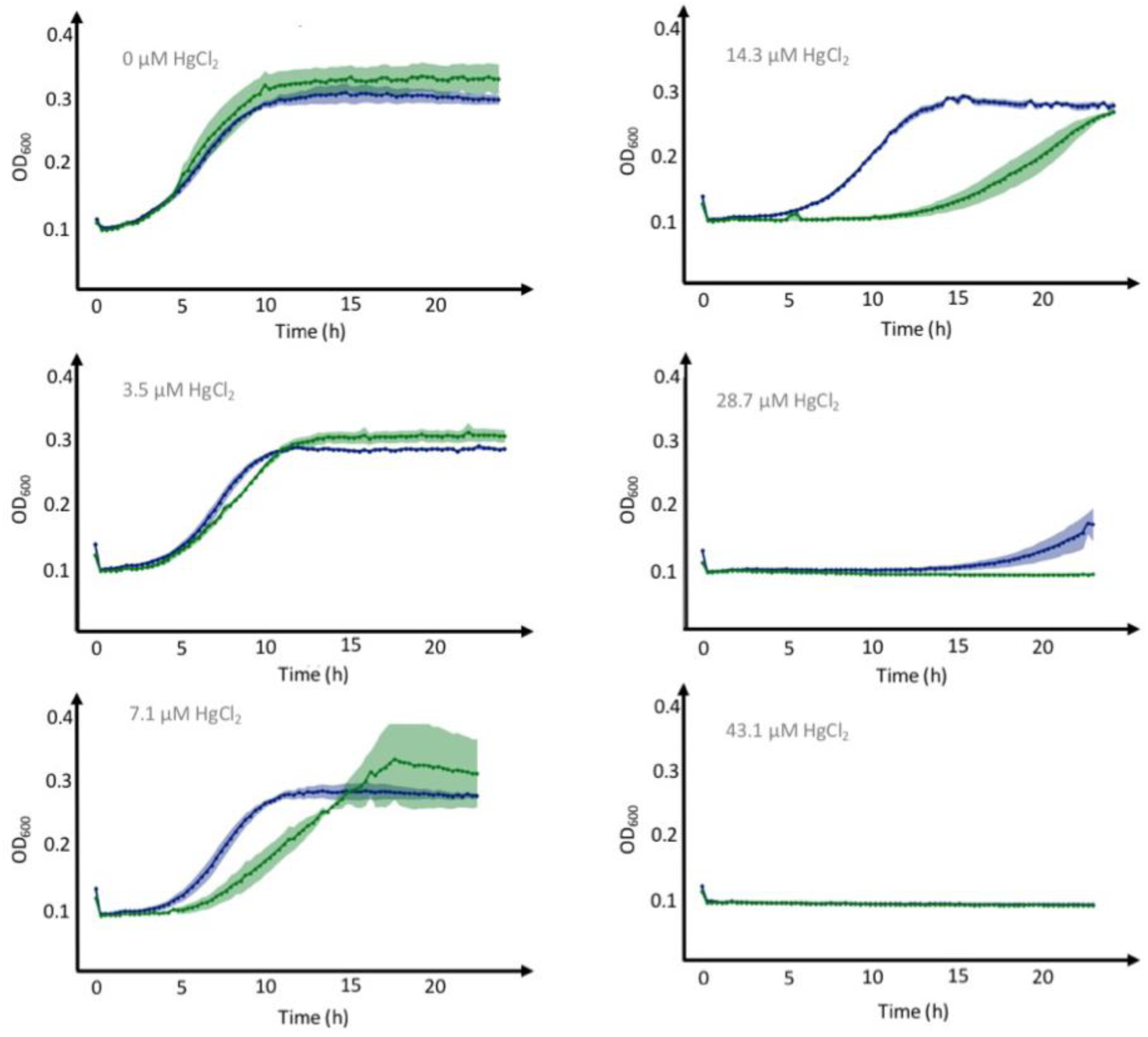
Improved mercury resistance in *Escherichia coli* DH10B containing p5343_UC57 (blue) versus only the empty vector pUC57 (green), grown in the presence of varying mercuric chloride concentrations.

**Fig 3:**
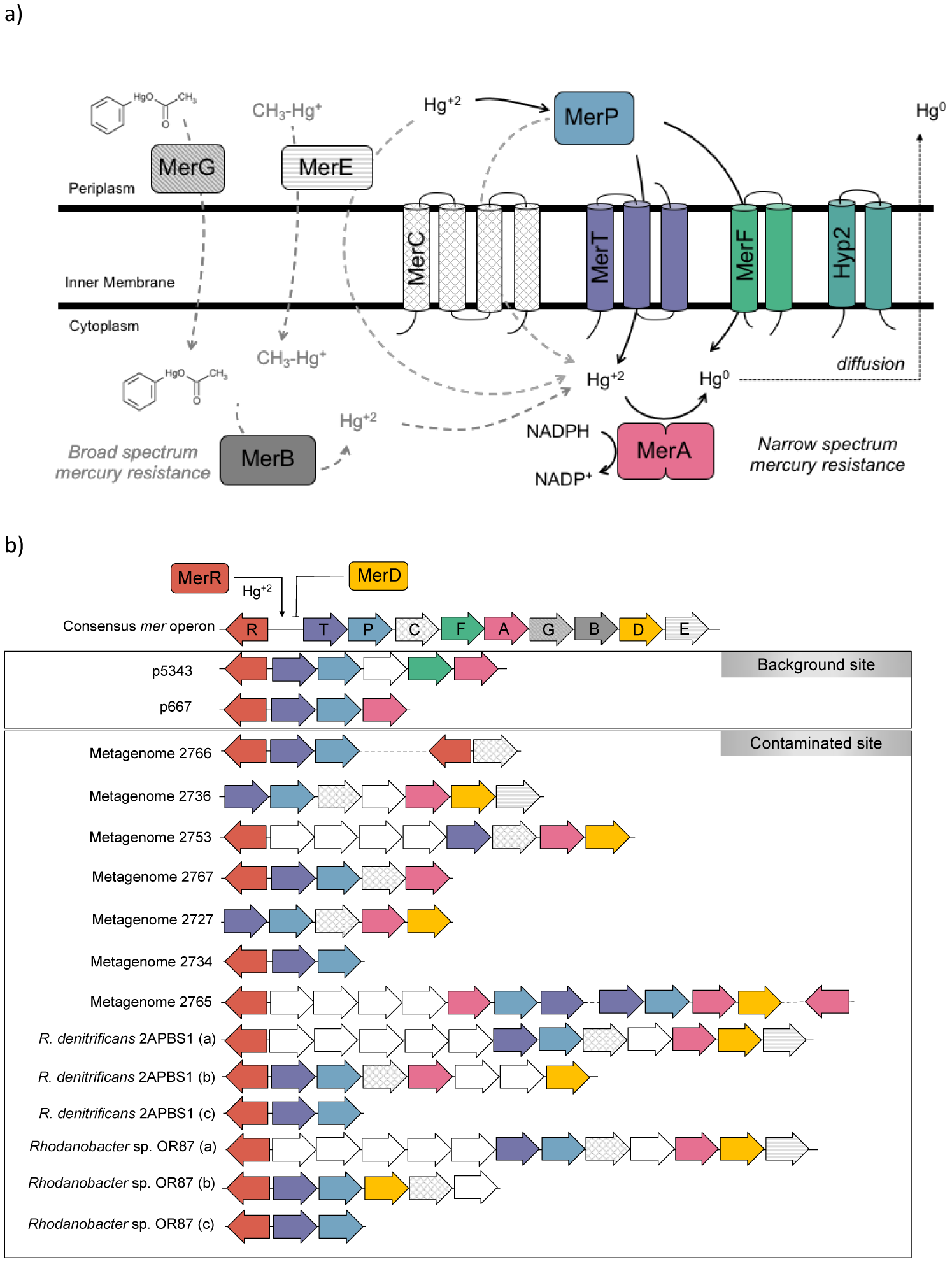
a) Schematic model of mercury resistance encoded by the *mer* operon, the genes absent on the plasmid p5343 are depicted in grey. b) The consensus *mer* operon (not drawn to scale) is provided for reference along with the putative *mer* operons from the background and contaminated sites as available from publishes sources^11, 14^.

Since elevated levels of mercury have been reported at the floodplain of East Fork Poplar Creek (contaminated site), which is close to the background site studied (23), the movement/flow of microbes between the two sites may be responsible for the presence of *mer* genes in a location with undetectable mercury. To evaluate this possibility, we used existing metagenome (12) and whole genome sequences (24) to compare the *mer* sequences from the contaminated site with those from the plasmidome of the background site. The *mer* operon structures differed between the background and contaminated sites (Fig 3b). Further, we performed phylogenetic analysis on the MerA sequences from the ORFRC along with known sequences available on NCBI for providing context. The overall pattern of MerA distribution (Fig 4) was similar to the 16S rRNA based distribution, as reported earlier (25-27). Notably, no inference about HGT can be drawn because the tree includes MerA sequences from plasmidome and metagenome analysis where the source microbe encoding the plasmid is not known. Increased diversity in mercury resistance genotypes has been reported in the presence of low mercury concentrations (28). We confirm the same observation – we find that while the homologs of MerA from the background site are distributed across several bacterial and archaeal phyla, those previously observed from the contaminated site cluster mostly with Gammaproteobacteria. Additionally, while we observed the presence of genes encoding both narrow- and broad-spectrum mercury resistance in contaminated sites, only those encoding narrow-spectrum resistance were found in the circular plasmids from background sites (Fig 3b). Thus, it is evident that the background sites have a diverse set of *merA* genes, which appear different from the metal-contaminated areas.

**Fig 4:**
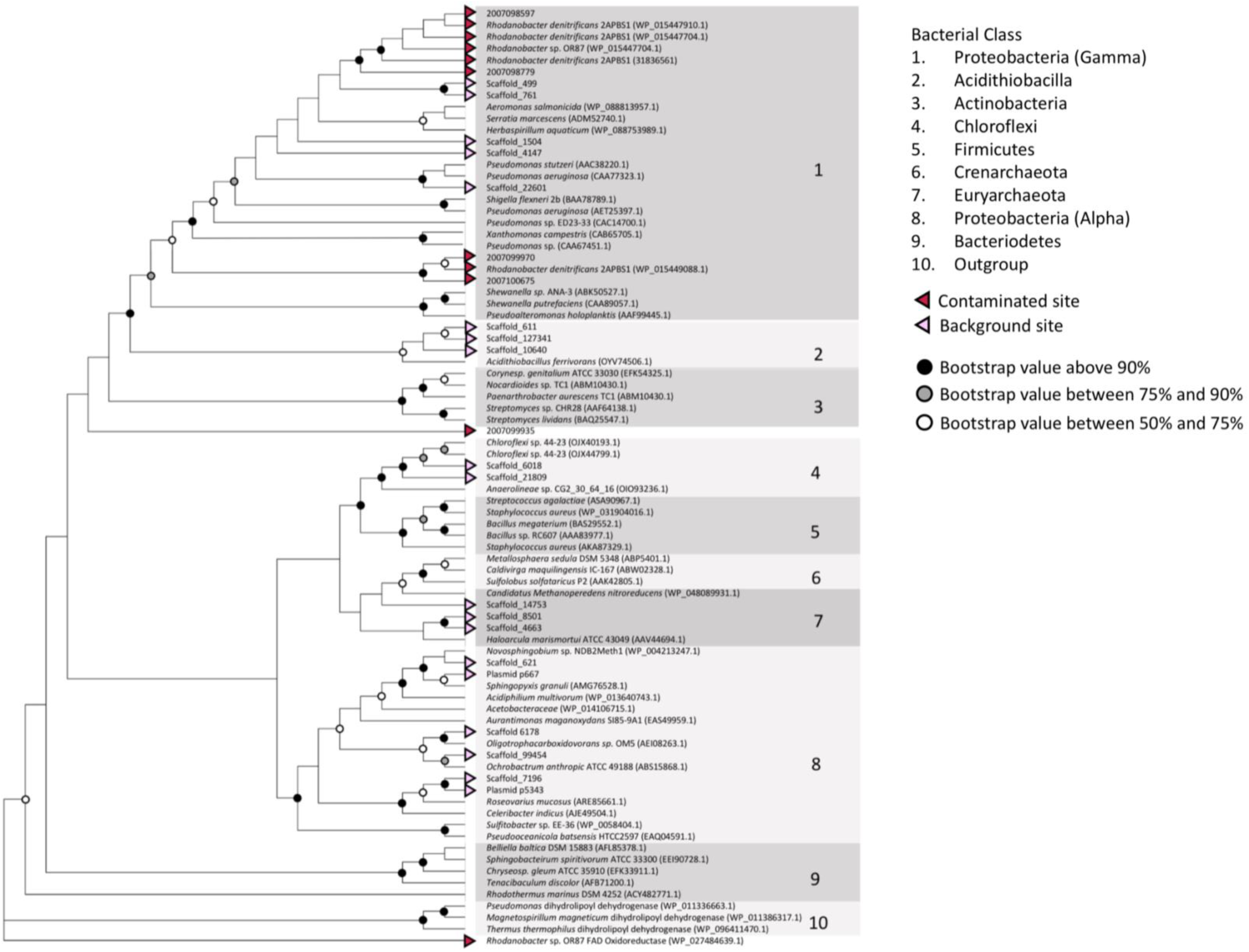
The evolutionary history based on MerA sequences was inferred by the maximum likelihood method using MEGA7^15^. The bootstrap consensus tree is inferred from 1000 replicates and bootstrap values (%) for well supported nodes are indicated. Coherent phyla are indicated to the right. The MerA sequences found in the background site are indicated by yellow triangles, while those reported previously from contaminated site are indicated by orange triangles.

Next, we designed an assay to test if the native plasmid-encoded metal resistance trait can be disseminated into other bacteria from this environment. Since this environment has very low cell density, natural transformation might be a prevalent HGT mechanism. Given that the ecological relevance of natural transformation may be better understood by studying bacteria and plasmids (DNA sequence is known to influence its uptake (29, 30)) native to this site, we used p5343 to test a suite of groundwater isolates for HGT of the plasmid-encoded mercury resistance trait. We find that the strains tested fall into three distinct categories (Fig 5). The first category includes the negatives *i.e.*, strains that do not show improved growth in mercury chloride in a plasmid-dependent manner. The second set included false positives *i.e.*, strains that showed improved plasmid-dependent growth in the presence of mercury chloride but were negative for colony PCR against *merA* gene. The third category included the positives *i.e.*, strains that show both improved growth in mercury chloride in a plasmid-dependent manner and were positive for colony PCR for the presence of *merA.* The positives included 17 strains belonging to six different genera and consisted of both Gram-positive (2 strains) and Gram-negative (15 strains) bacteria. Amongst these, natural competence had never been reported in the genera *Arthrobacter, Dermacoccus, Acidovorax*, and *Cupriavidus*. Thus, the screen identified 17 strains that acquired mercury resistance and likely took up the plasmid via natural transformation.

**Fig 5:**
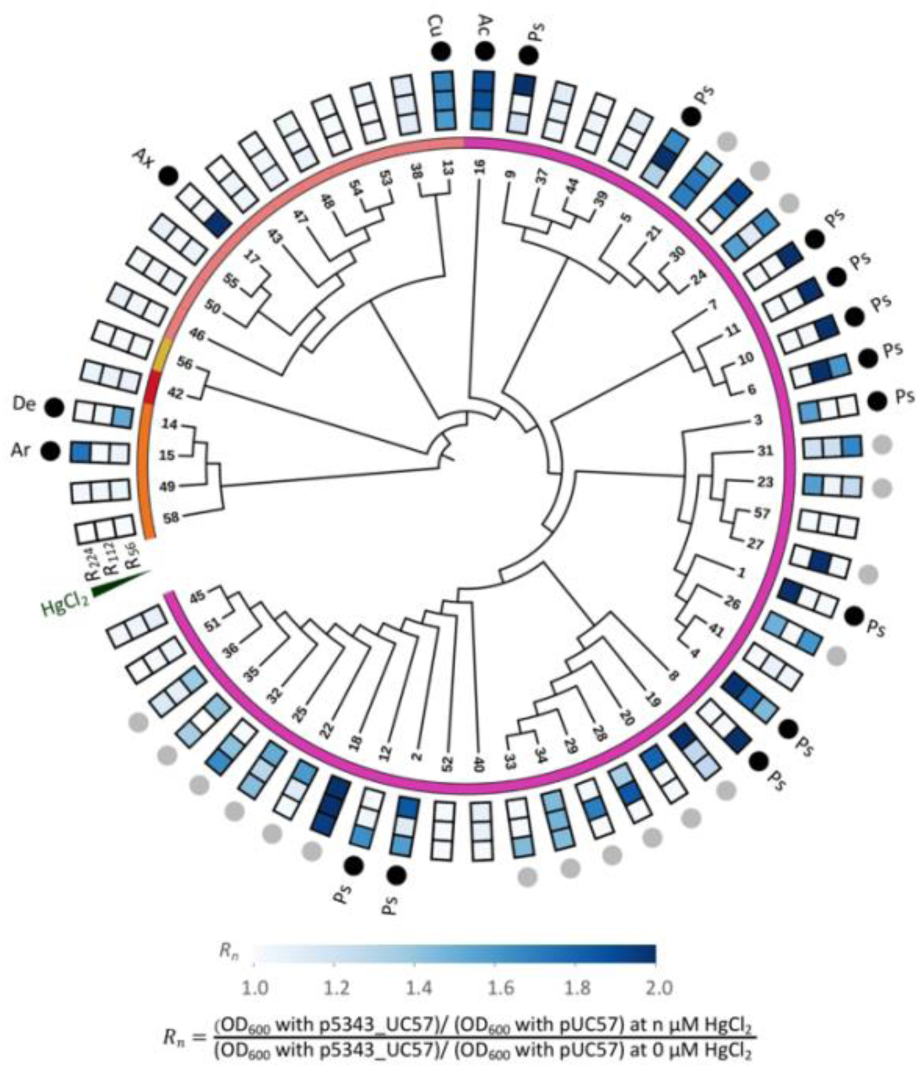
Results of the HGT assay used to screen environmental isolates for uptake of a plasmid-encoded trait via natural transformation. The 58 bacterial isolates are depicted on a 16S rRNA sequence-based phylogenetic tree (constructed using Maximum Likelihood). The colored bars indicate different bacterial classes, namely, orange is Actinobacteria, red is Cytophagia (Bacteriodetes phylum), yellow is Alphaproteobacteria, peach is Betaproteobacteria and pink is Gammaproteobacteria. The heat map plots optical density in the presence of p5343_UC57 compared to pUC57 in the presence of different mercury chloride concentrations normalized by the no mercury chloride control. Only the strains showing improved plasmid-based growth in the presence of mercury chloride were subjected to *merA* based colony PCR. The strains positive for colony PCR are indicated in black circles while those negative are depicted in grey circles. The genus of the strains positive for natural transformation as depicted on the figure are *Arthrobacter* (Ar), *Dermacoccus* (De), *Acidovorax* (Ax), *Cupriavidus* (Cu), *Acinetobacter* (Ac) and *Pseudomonas* (Ps). Figure constructed using ITOL^46^.

For further analysis, two *Pseudomonas* strains (from the 17 positives), along with model strain *E. coli* DH10B, were subjected to a more comprehensive natural transformation assay with replicates. This involved incubation of strains with plasmids for natural transformation, followed by growth under mercury stress for two serial transfers. The *Pseudomonas* strains 5 and 12, along with DH10B demonstrated plasmid-dependent improved growth in the presence of mercury chloride over the two serial growth regimes (Fig 6). For strain 12, the improved growth was more evident at a lower mercury chloride concentration than at higher concentrations. Following the second growth regime, all three strains were positive for p5343 encoded *merA* via qPCR after the assay while being negative before the assay. Thus, we confirm HGT and functionality of the p5343 encoded *mer* operon in a model bacterium and field-relevant environmental bacteria.

**Fig 6:**
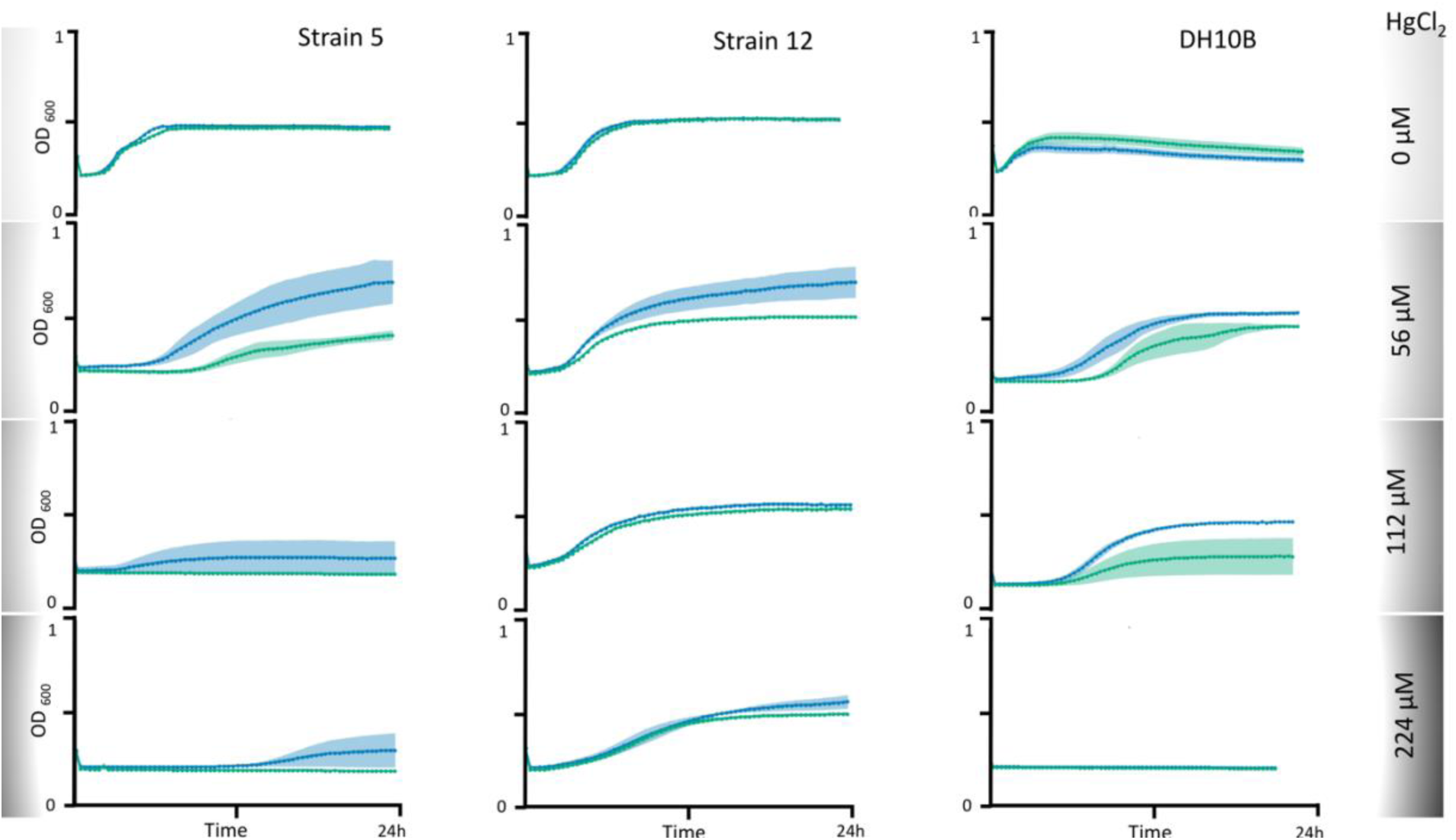
Confirmation of HGT in strains 5, 12 and DH10B. Growth curves from second growth regime in the presence of varying mercury chloride concentrations are depicted when strains 5, 12 and DH10B were naturally transformed with plasmid p5343_UC57 (blue) and pUC57 (green). Assay design provided in Figure S2.

## Discussion

The presence of both background and metal-contaminated sites at ORFRC makes it an excellent system to study the prevalence and dissemination of metal-resistance determinants in microbial communities. Analysis of mercury reductases found in the contaminated and background sites revealed a significant decrease in the gene diversity in the former compared to the latter. This might be due to better functioning of a particular homolog(s) of *merA* in the presence of high levels of heavy metals (particularly mercury) and/or the inability of certain phyla (encoding diverse *merA* homologs) to survive in such conditions, given the reduced taxonomic diversity in contaminated groundwaters. Alternatively, it is possible that the contaminated site sequences available do not capture the entire *merA* diversity and a targeted plasmidome analysis from the contaminated groundwater can shed light on this matter. Given that the background site is off-gradient from the contaminant plume and that the mercury resistance determinants based on the phylogenetic tree vary between the two sites, there is a reduced likelihood of bacterial carry-over from the contaminated to the background site. The high abundance of the *mer*-encoding plasmid in the absence of detectable mercury in the background site could indicate very effective mercury reduction and volatilization of any trace amounts of mercury that leached into the groundwater, by the resident microbiome community. A study demonstrated that plasmid persistence could be attributed to compensatory adaptation, along with brief periods of positive selection (31), which might be the most plausible explanation for the persistence of metal resistance gene(s) in plasmids from the background site.

The spread of mercury resistance via conjugation has been well-documented in soil microbial communities (32, 33). In contrast, since aquatic environments are typically characterized by low cell densities and availability of exogenous dissolved DNA (1.7 – 88 μg per liter (34)), we hypothesized that natural transformation could be a HGT mechanism in these systems. Based on our screen studying the dissemination of mercury resistance determinants via natural transformation, several environmental strains tested were either negatives or false positives. This could be attributed to the strains not being naturally competent in the conditions tested or were competent but not able to maintain/express the *mer* genes encoded on the plasmid or the *mer* provided resistance at mercury concentrations lower than that tested. The presence of *relE* toxin gene on p5343 plasmid could also influence the retention and expression of the plasmid in certain strains, limiting it to strains that carry the corresponding anti-toxin gene. Interestingly, we identify 17 strains that acquired an environmental plasmid-encoded trait via natural transformation. These comprised of 12 *Pseudomonas* strains, which was notable because prior work showed that linear dsDNA was effective whereas ssDNA or plasmid DNA did not work for natural transformation in *Pseudomonas* (35). In fact, in strains known to be naturally competent, genomic DNA is typically preferred over circular plasmid sequences (35, 36), a trait likely attributed to the requirement of linearizing the circular plasmid prior to uptake. Thus, it is significant that employing a circular native plasmid, we demonstrate HGT via natural transformation into environmental bacteria, indicating this could be a viable route for plasmid dissemination in groundwater.

Successful HGTs frequently occur between closely related organisms (37), and the compositional similarity between the donor and the recipient genomes promotes homologous recombination leading to DNA acquisition from close relatives. Interestingly, although all the p5343 encoded genes are closest to Alphaproteobacteria, the strains that were transformed belong to Betaproteobacteria, Gammaproteobacteria and Acinetobacteria. Amongst the 58 strains, only one belonged to Alphaproteobacteria, and testing more strains from this class might result in more positives. Natural competence is a transient physiological cell state which allows DNA uptake under specific conditions (9) (an exception being *Neisseria gonorrhoeae* where competence is constitutive (38)). Thus, the ideal set of conditions for competence are strain-specific and the screen can be further optimized by varying plasmid incubation times, the amount of plasmid added, the cell density at which the plasmid was added, changes in media constituents, and testing other mercury concentrations. Additionally, in its native state, the p5343 may be methylated, and further experiments with different methylated versions of p5343 plasmid might enable HGT into additional bacterial phyla from this environment.

Overall, this study reveals that the highly abundant *mer-*encoding plasmids are functional in providing mercury resistance and capable of being horizontally transferred into relevant bacterial isolates from the ORFRC site. We demonstrate that natural transformation facilitates the inter-phyla transfer of genetic elements, suggesting that the transient presence of plasmid DNA in close vicinity may be sufficient for HGT to occur in the groundwater communities. This suggests that the microbial community studied is likely robust in tolerating low stresses and possesses a latent ability to swiftly adapt to changes in the environmental stress levels using natural transformation.

## Materials and Methods

### Plasmid p5343

The identification and annotation of plasmid p5343 has been previously described (14). The 8 kb p5343 was synthesized by Genscript (Piscataway, NJ). To propagate the synthesized DNA in the model bacterium *E. coli*, p5343 was cloned into the vector pUC57 resulting in plasmid p5343_UC57 (plasmid and sequence available via Addgene 126645).

### Analysis of mer genes

For the phylogenetic analysis of MerA, amino acid sequences were obtained from the plasmidome (14) of the background site along with the metagenome (11, 24) and whole genome sequences (11, 24) of the contaminated site at ORFRC (given the lack of availability of plasmidome data from the contaminated site). In addition, we also added publicly available MerA sequences from NCBI (accession numbers provided on the phylogenetic tree). The evolutionary history based on MerA sequences was inferred using MEGA7 (39). The MerA sequences were aligned using Muscle (40). The alignment was manually curated, and all the MerA sequences were cropped to a common length of 515 amino acids. The JTT matrix-based model (41) was used to construct maximum likelihood trees with 1000 bootstrap replicates (42).

### Mercury Resistance

The plasmids pUC57 and p5343_UC57 were transformed into *E. coli* strain DH10B strain separately. Five freshly transformed colonies were picked for each plasmid and overnight cultures were prepared at 37 °C. To compare the growth of DH10B transformed with p5343_UC57 and pUC57, 5 μl of pre-culture (final OD_600_ of 0.1) was inoculated into 95 μl of LB medium with carbenicillin (100 µg/ ml final concentration) in a 96 well transparent flat bottom tissue culture plate (Corning, Falcon ®, catalog number: 353003) in the presence of 0, 3.5, 7.1, 14.3, 28.7 and 43.1 μM final mercury chloride concentrations. The plate was sealed with BreathEasy seals (E&K Scientific, Santa Clara, CA, USA) and grown at 37 °C in a Tecan F200 microtiter plate reader (Tecan Group Ltd., Männedorf, Switzerland) with shaking, measuring the optical density at a wavelength of 600 nm at 20 min intervals.

### HGT via High-Throughput Screen

Isolation, arraying, and recovery of the bacterial strains from ORFRC was done as previously described (43). We re-arrayed and screened 58 previously described bacterial strains (43). The strains are numbered 1 through 58 for ease of representation in figures and text (details in Table S1). The design of the assay to screen for HGT in the bacterial isolates is described in Fig S1. The arrayed isolates were recovered in a 96 deep-well plate (Costar, Thermo Fisher Scientific, Waltham, MA, USA) by addition of 1 ml of appropriate growth (either LB or R2A) media. The 96 deep-well plate was grown in Multitron plate shaker/incubator (Infors, Bottmingen, Switzerland) at 30 °C with shaking at 700 rpm, overnight. The 96 well plate was centrifuged at 4000 rpm for 10 min in Eppendorf 5810R centrifuge and the pellets were resuspended in 1 ml fresh growth media. Approximately 250 μl of each culture was added to individual wells of two 96-well deep well plates. Approximately 5 μl of 50 ng/μl p5343_UC57 was added to each well of one plate while pUC57 was added to each well of the second plate. Both the plates were incubated overnight at 30 °C without shaking to enable natural transformation. Since improved competence has been reported in cells which are stressed (8), after 24 h, 40 μl of the un-diluted overnight cultures from the two 96 well plates were moved to two 384-well flat bottom transparent plates (Corning, catalog number: 353003) using a Biomek FxP (Beckman Coulter) liquid handling robot. To each quadrant of the 384-well plate 40 μl of appropriate media with 0, 112, 224, 448 μM mercury chloride concentration was added such that final concentrations were 0, 56, 112 and 224 μM mercury chloride. The plates were sealed using BreathEasy seals (E&K Scientific, Santa Clara, CA, USA). The growth of both 384 well plates was monitored over 24 h at 30 °C in Tecan F200 microtiter plate reader with shaking, measuring the optical density at a wavelength of 600 nm at 20 min intervals (growth data provided in Table S2). Since certain strains might already encode a native *merA*, we look for improved mercury resistance in presence of p5343_UC57 versus pUC57. The strains with plasmid-dependent improved growth in the presence of mercury chloride were subjected to colony PCR using primers 5’-cacaccgcccaaaagtctat-3’ and 5’-cagagctggcacagatgatg-3’ designed to amplify the *merA* gene. All the cultures tested were confirmed to be negative for *merA* by colony PCR before the assay. For all strains that were positive for *merA* colony PCR post assay, an additional round of 16S rRNA sequencing was done to confirm the identity of the strains.

To obtain a phylogenetic tree depicting the 16S rRNA sequences of all 58 strains tested, the 16S rRNA sequences of these strains were aligned using Muscle (40). The alignment was manually curated, and evolutionary history was inferred using the Minimum Evolution (ME) method (44). The evolutionary distances were computed using the Maximum Composite Likelihood method (45) and are in the units of the number of base substitutions per site. The ME tree was searched using the Close-Neighbor-Interchange (CNI) algorithm at a search level of 1. The Neighbor-joining algorithm (46) was used to generate the initial tree. Evolutionary analyses were conducted in MEGA7 (39).

### Confirmation of HGT in Selected Strains

The design of the natural transformation assay for confirmation of HGT in selected strains is depicted in Fig S2. Frozen stocks of the ORFRC isolate strain 5 (*Pseudomonas* FW305-3-2-15-A-LB2) and strain 12 (*Pseudomonas* FW305-20), along with *Escherichia coli* DH10B were recovered overnight in culture tubes with 5 ml LB media grown at 30 °C. The cultures were centrifuged and fresh 2.5 ml LB media was added to the pellet, followed by resuspension. About 250 μl of each culture was added to individual wells of a 96 deep-well plate (Costar, Thermo Fisher Scientific). To each well, 5 μl of the appropriate plasmid (p5343_UC57 or pUC57) at a concentration of 50 ng/μl was added. The plate was incubated at 30 °C without shaking to enable natural transformation. Post 24 h, 50 μl of the un-diluted overnight cultures with plasmids from each well was transferred to four wells of a 96 well transparent flat bottom tissue culture plate (Corning, Falcon®, catalog number: 353003) and 50 μl of LB media with 2X mercury chloride concentrations was added to each well such that the final concentrations were 0, 56, 112, and 224 μM mercury chloride. The growth was monitored over 24 h at 30 °C in Tecan F200 microtiter plate reader with shaking, measuring the optical density at a wavelength of 600 nm at 20 min intervals after which 5 µl from each plate was added to a fresh 96 well plate. To perform a second growth regime in the presence of mercury chloride, to each well 45 ul of LB and 50 ul of LB with 2X mercury chloride was added (to reach the same final mercury chloride concentrations) and the growth was monitored over 24 h at 30 °C in Tecan F200 microtiter plate reader. The 24 h growth data was analyzed to see if the strains with p5343_UC57 had an advantage over pUC57 when grown in the presence of mercury chloride. In addition, all strains were subjected to quantitative PCR with primers 5’-cgtccttgtcgaaggtttgt-3’ and 5’-atagacttttgggcggtgtg-3’ designed to amplify p5343-encoded *merA* both before (using glycerol recovered overnights) and after the assay (qPCR raw data provided in Table S3). The qPCR was performed using EvaGreen Real-time PCR (Biotium, Hayward, USA) according to the manufacturer’s recommendations using 1µl EvaGreen Real-time PCR (Biotium, Hayward, USA), 1 µl of cell culture, 12.5 µl of Q5 High-Fidelity 2X Master Mix (New England Biolabs, Ipswich, MA, USA), 1.25 µl each of 10 µM forward and reverse primers, and 9 µl of nuclease-free water.

## Funding Sources

This work was part of the ENIGMA-Ecosystems and Networks Integrated with Genes and Molecular Assemblies (http://enigma.lbl.gov), a Science Focus Area Program at Lawrence Berkeley National Laboratory and is supported by the U.S. Department of Energy, Office of Science, Office of Biological & Environmental Research under contract number DE-AC02-05CH11231 between Lawrence Berkeley National Laboratory and the U. S. Department of Energy. The funders had no role in study design, data collection and interpretation, or the decision to submit the work for publication. The United States Government retains and the publisher, by accepting the article for publication, acknowledges that the United States Government retains a non-exclusive, paid-up, irrevocable, world-wide license to publish or reproduce the published form of this manuscript, or allow others to do so, for United States Government purposes.

## Acknowledgements

We thank Megan Garber (LBNL) and Nurgul Kaplan (LBNL) for help with liquid handling systems. We thank Han Zhang (LBNL) for his help with figures. We thank John-Marc Chandonia (LBNL) for help with the environmental strains sequence data tracking. The funders had no role in study design, data collection and interpretation, or the decision to submit the work for publication.

## Conflict of interest

Authors do not have any conflict of interest.

## Supplementary Information

### List of Supplementary Tables

Table S1 : Details of the strain names, isolation conditions, taxonomic assignments, and 16S rRNA sequence of 58 strains subjected to HGT assay screen.

Table S2 : The raw and normalized (to no mercury control) ratios of optical density measured at 600nm in presence of p5343_UC57 compared to pUC57 after 24h of growth in the presence of different mercury concentrations.

Table S3 : Results from qPCR depicting quantitation cycle (Cq) values indicating the presence of *merA* gene in strains 5, 12 and DH10B post the second growth regime in mercury chloride in the comprehensive natural transformation assay. The lower the Cq value, the higher initial copy number of the target gene.

### List of Supplementary Figures

Fig S1 : Design of the HGT assay to screen environmental bacterial isolates for uptake of a plasmid-encoded trait via natural transformation.

Fig S2: Design of the natural transformation assay for the confirmation of HGT in the strains 5, 12 and DH10B (indicated by different shades of grey) in replicates.

